# Fine-tuning *RIPENING INHIBITOR* (*RIN)* expression by introducing allelic mutations in its promoter using CRISPR/Cas9 multiplex editing

**DOI:** 10.1101/2025.07.14.664838

**Authors:** Jiaqi Zhou, Chiu-Ling Yang, Diane M. Beckles

## Abstract

Tomato is one of the most produced and consumed vegetables globally due to its nutritional benefits, sensory characteristics, and cultural importance. However, tomato fruit has a short shelf-life, which can be extended by postharvest techniques, but often at the expense of fruit quality, leading to consumer dissatisfaction. To address this challenge, we modified the upstream regulatory regions of Ripening inhibitor (RIN), a master regulator of tomato fruit ripening, utilizing a CRISPR/Cas9 multiplex system. This approach enabled the creation of a population of tomato fruit with mutations of varying severity, which could have far-reaching effects on the RIN-induced gene regulatory network in tomato fruit, leading to downstream changes in fruit traits. We have generated 264 first-generation (T_0_) transgenic lines of RIN promoter mutants with diverse genetic lesions and RIN transcriptional levels. Our study revealed a non-linear relationship between promoter mutations and gene expression, highlighting the potential roles of certain types of mutations in regulating RIN transcription. Future work will focus on evaluating fruit traits from mutants with pronounced changes in *RIN* expression, as well as performing transcriptomic analysis to explore the mechanisms underlying fruit quality modifications due to genome editing.

## INTRODUCTION

Tomato is a valuable vegetable crop but is perishable, and thus has a limited shelf-life. It is also an important functional genomics model for studying the regulation of fleshy fruit ripening, which is complex. Ripening is governed by many Transcriptional Factors (TFs) which orchestrate the expression of other TFs and downstream genes as part of multiple, but highly integrated ripening regulatory networks (Tonutti et al., 2023). The MADS-box RIPENING INHIBITOR (RIN) directly binds to many downstream TFs and genes, turning them on/off during fruit ripening (Fujisawa et al., 2013). The naturally occurring tomato *rin* mutant, was discovered in the 1900s, and its allele has been incorporated into tomato breeding programs to extend fruit shelf-life because it slows ripening. Recent studies revealed that *rin* is a gain-of-function mutation, and a CRISPR/Cas9 null mutant *Rin* was capable of initiating fruit ripening but disrupted ripening phenotypes (Ito et al., 2017; Li et al., 2020; Vrebalov et al., 2002). Due to severe disruptions in *RIN* expression, many of the desirable quality traits associated with fruit ripening are destroyed (Ito et al., 2017; Li et al., 2020; Vrebalov et al., 2002).

Editing gene regulatory regions instead of coding regions may provide an opportunity to fine-tune gene expression, therefore leading to a range of subtle phenotypic changes (Albornoz et al., 2022; Huang et al., 2020). This work focuses on *RIN* transcriptional regulation. *RIN* gene promoter regions, upstream of the transcription start site, contain a series of *Cis*-Regulatory Elements (CREs), i.e., specific DNA sequence motifs to which TFs bind, to drive or suppress *RIN* gene expression (Schwarzer & Spitz, 2014). CRE mutations can destroy TF binding sites, introduce additional copies of a CRE, or create a novel CRE to allow new TFs to bind, thereby altering the transcriptional levels of its cognate gene (Schwarzer & Spitz, 2014). Analysis of the *RIN* promoter (Fig. 1A) indicates that there are (a) two CArG motifs to which RIN binds (Bemer et al., 2012; Fujisawa et al., 2013), (b) two G-box motifs that the TF Elongated Hypocotyl 5 (HY5) binds to during tomato fruit ripening (Wang et al., 2021), and, (c) numerous putative CREs to which diverse TFs may bind in response to changes in developmental stages, environmental conditions, and hormones (Lescot et al., 2002). The CArG motifs are of interest because RIN protein binding may create an autoregulatory loop that intensifies the regulation of *RIN* gene transcriptional levels, with consequences for downstream TFs and genes.

**Figure 1.**
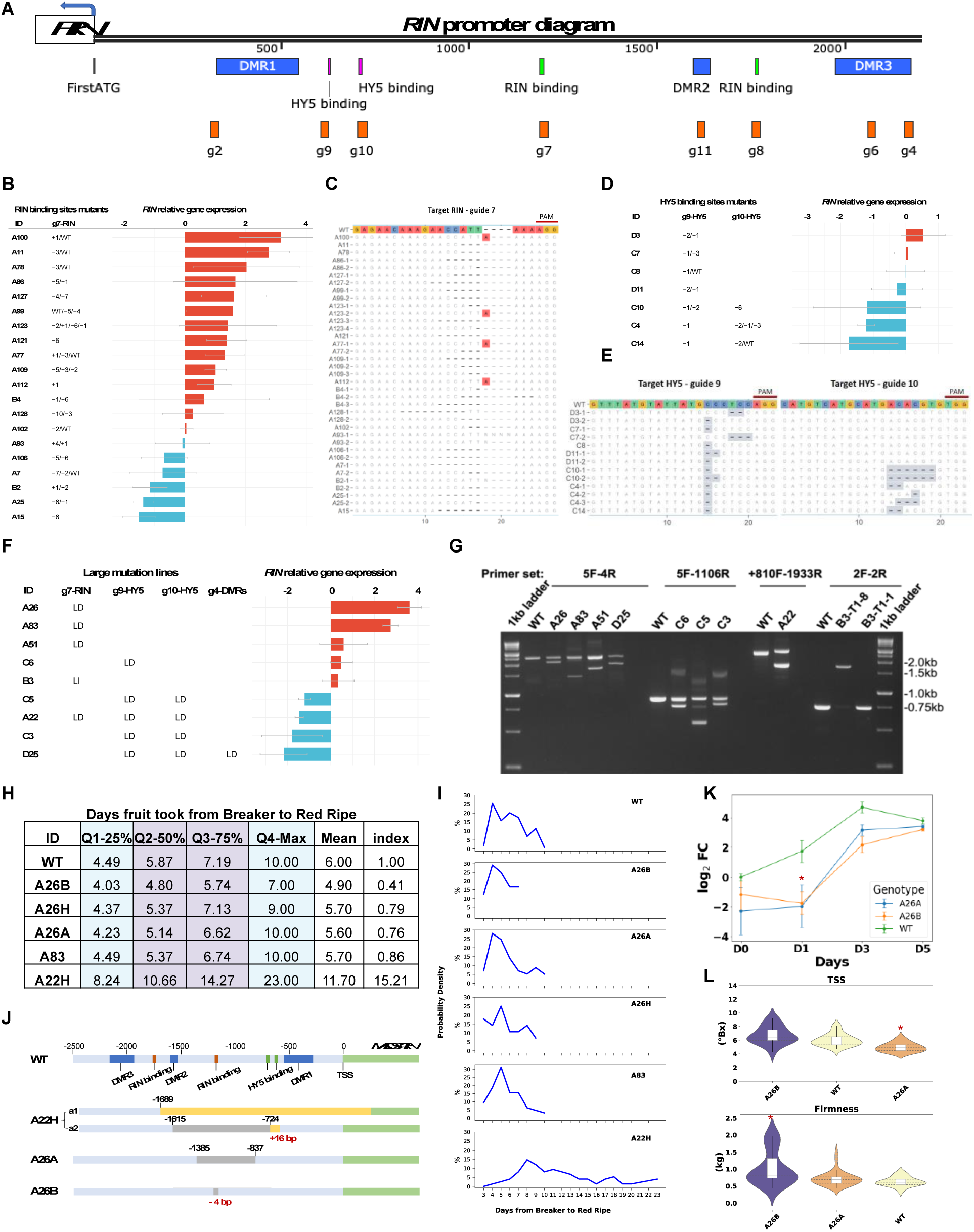
*RIN* promoter allelic mutants using CRISPR/Cas9. **(A)** Analysis of the *RIN* promoter, arbitrarily defined as the 2-3 kb upstream of the transcriptional start site. There are three DMRs, targeted by gRNAs -g2, g6, and g4, respectively; two HY5 binding sites targeted by g9 and g10, and two RIN binding sites targeted by g7 and g8. The gRNA sequences are in Table S1. A summary of the mutations, their location, types, and levels are as follows: **(B)** and **(C)** a RIN binding site, **(D)** and **(E)** HY5 binding sites, **(F)** and **(G)** large mutation, i.e., large insertion (LI) or large deletion (LD) at a single RIN binding site, two HY5 binding sites, or DMRs. *RIN* relative gene expression in tomato fruit at red ripe was normalized to the reference gene *SlCAC*, using WT red-ripe fruit as the control, and primers used are in Table S4. The log_2_ fold change was presented as the bar in **(B, D, F)**. The red bars show *RIN* upregulation-log_2_ fold change > 0, and the blue bars show *RIN* downregulation relative to the WT-log_2_ fold change < 0. The sequence of the mutated alleles for **(B)** and **(D)** is shown as **(C)** and **(E**). In **(C)** and **(E)**, the color of the nucleotide backgrounds indicates mutation types; white indicates no change compared to the WT, grey shows deletions, and red regions indicate insertions. There are multiple alleles in some mutants, which are presented in individual rows. **(G)** A 1% (w/v) agarose gel image showing one or multiple alleles in the large mutation lines, using WT as the control. The primer pairs used to amplify the bands are indicated 5F-4R etc. (details are in Table S3). **(H)** Quartile statistic and index of ripening days in selected T_1_ edited lines. Data shows the number of days fruit took to go from Breaker to Red Ripe when ripened on-the-vine. Five mutants and the WT are included. A ripening index for each genotype was computed by multiplying all values from Q1-Q4 and dividing the result by the corresponding value in the WT. **(I)** Trends of fruit on-the-vine ripening over time based on data generated from **(H).** The x-axis represents the number of days from Breaker to Red Ripe, and the y-axis shows the probability density, indicating the proportion of fruit within the population. The purple shaded area highlights the interquartile range (Q2 to Q3), and the mean number of days is marked within this range. **(J)** Genotypes ‘A22H’, ‘A26A’ and ‘A26B’ showing the locations of introduced mutations in the *RIN* gene promoter. **(K)** *RIN* relative gene expression during on-the-vine ripening. The y-axis shows log_2_ fold change of *RIN*, using the WT at Breaker stage (D0) as the control. The x-axis represents days post-breaker (0, 1, 3, 5 days). **(L)** Fruit quality traits, including total soluble solids (TSS), and firmness (kg). Fruit were harvested at the breaker stage and stored at 20°C for one week. Dunnett’s pairwise comparison was performed using WT as the control. Asterisks ‘*’ indicate statistically significant differences (*p* < 0.05).

RIN binding sites in the gene promoter are frequently close to differentially methylated regions (DMRs), with RIN binding usually occurring at hypomethylation sites (Zhong et al., 2013). DMRs vary in DNA methylation status due to developmental or environmental cues, and may influence transcription by modulating the chromatin accessibility of nearby TFs (Henderson & Jacobsen, 2007). In the *RIN* promoter, DMRs have been identified that are hypermethylated in immature green tomato fruit but then, are demethylated in ripe fruit (Lang et al., 2017). Global DNA demethylation is crucial for tomato fruit ripening and for RIN binding its downstream target TFs and genes. The tomato *dml2-3* mutant, which lacks a key DNA demethylase, shows genome-wide hypermethylation and disrupted fruit ripening (Niu et al., 2025). Ectopic expression of *RIN* in the *dml2-3* background partially restores the ripening phenotype, possibly by restoring demethylation at the RIN binding sites (Niu et al., 2025). However, it remains unclear whether changes in DMRs within the *RIN* promoter, rather than genome-wide methylation shifts, are sufficient to influence *RIN* transcription and fruit ripening.

Evidence suggests that editing multiple CREs in the promoter of key regulatory genes in tomato, can lead to subtle alterations in the mRNA levels of the edited gene (Rodríguez-Leal et al., 2017). Therefore, differentially editing the *RIN* promoter using multiplex CRISPR/Cas9 could fine-tune *RIN* mRNA and protein levels. In addition to the known TF binding sites, recent findings reveal that SlWOX13, a member of the WOX family and a newly identified ripening enhancer, also binds the *RIN* promoter adjacent to the HY5 binding site (Jiang et al., 2023). This finding illustrates the complexity of the RIN regulatory landscape, where multiple TFs converge on the *RIN* promoter to coordinate its transcription during ripening. Therefore, subtle changes in *RIN* gene promoter sequence, could lead to a cascade of nuanced alterations in ripening-related gene expression patterns that would affect ripening rates and fruit quality. Taken together, this work is expected to answer whether mutations in the *RIN* gene promoter can affect its transcription and create novel ripening phenotypes.

## RESULTS & DISCUSSION

We designed eight 20 bp-gRNAs spanning the *RIN* promoter, targeting each regulatory site (Fig. 1A). There were four sets of gRNAs combinations assembled, to target regulatory regions, individually or concurrently, i.e., (a) two gRNAs targeting the identified RIN self-binding sites (the CArG motif) (Bemer et al., 2012) - g7 and g8, (b) two gRNAs targeting the HY5 binding (Wang et al., 2021) sites - g9 and g10, (c) four gRNAs on the DMRs (Fig. S1) (Lang et al., 2017) - g2, g11, g4, and g6, and (d) six gRNAs targeting the RIN and HY5 binding sites and the DMRs together - g2, g9, g10, g7, g8, and g4 (Fig. S2). Four CRISPR/Cas9 constructs were assembled and introduced into tomato cotyledons using *Agrobacterium*-mediated plant transformation. We generated 264 independent first-generation (T_0_) transgenic lines. The genotyping results indicated that 24 lines had large insertions or deletions within the *RIN* promoter, ranging from hundreds to thousands of base pairs, sometimes spanning multiple CREs. Many lines had small indels at one or multiple sites; for mutants with small indels at a single site, 51 and 29 were at the RIN and HY5 binding sites, and 14 were at the DMRs. Only four mutants had small indels at both the HY5 sites and DMRs. The sequencing results showed that most T_0_ lines were heterozygous, containing at least one wild-type (WT) allele, or exhibited biallelic or chimeric patterns in the targeted regions. Only a few lines, such as ‘A121’, ‘A112’, ‘A15’, and ‘C14’, had homozygous alleles (Figs. 1B-E, S3).

To evaluate the effect of these promoter mutations on *RIN* transcription during fruit ripening, the relative expression of *RIN* in mutant fruit was compared to control fruit (WT) using RT-qPCR. Tissue culture plantlets were transferred into a greenhouse and T_0_ lines were selected for fruit tissue sampling. Only red-ripe fruit was assessed, as *RIN* expression is stable at this developmental timeframe in WT fruit (Li et al., 2020; Shinozaki et al., 2018). We observed a range of *RIN* expression levels among the T_0_ mutants, and ranked the lines accordingly based on their *RIN* expression (Fig. 1).

### RIN binding sites

T_0_ lines with mutations at the *RIN* CRE by g7 (Fig. 1B) had both positive and negative effects on *RIN* expression. However, no edits were associated with g8 when over a hundred lines were screened. This is possibly due to g8’s low editing efficiency or limited chromatin accessibility at its target site. *RIN* expression in ‘A100’ was 8.9-fold higher, while in ‘A15’, it was 2.9-fold lower than that in the WT. By examining the mutation location and expression levels (Fig. 1B, C), the data suggested (i) a single adenine insertion may underlie the higher *RIN* expression observed in ‘A100’, ‘A123’, ‘A77’ and ‘A112’, (ii) both ‘A11’ and ‘A78’ had a 3-bp deletion and are heterozygous to WT, with an upregulation of *RIN*, (iii) ‘A15’ and ‘A121’ had contradictory *RIN* expression pattern but the same type of promoter mutation; somaclonal variation, induced by tissue culture, may differentially influence *RIN* expression (Bairu et al., 2011).

### HY5 binding sites

Simultaneous mutations at both HY5 sites by g9 and g10 led to a downregulation of *RIN* expression (‘C10’, ‘C4’, ‘C14’). Four of the seven lines were reduced in *RIN* expression, one was unchanged, and the other two lines were slightly upregulated (Fig. 1D). It is of note that the location at which the editing occurs is important to transcription; an inverse expression pattern was observed in ‘D3’ and ‘D11’ – these individuals are heterozygous for a 2- bp and a 1-bp deletion (Fig. 1D), but the deletions are at different locations within the g9 target region (Fig. 1E). This variation potentially leads to divergent *RIN* expression (Fig. 1D).

### Large mutations

Two heterozygous large deletions T_0_ lines, ‘A26’ and ‘A83’, had upregulated *RIN* expression (Fig. 1F, G). ‘A26’ is a biallelic mutant with different deletions at a RIN binding site. The ‘A26’ T_1_ generation included the heterozygote ‘A26H’, and two homozygous genotypes, i.e., ‘A26B’ with a 4-bp deletion, and ‘A26A’ with a 321-bp deletion (Fig. 1J).

However, the three ‘A26’ T_1_ genotypes did not show the same pattern as their T_0_ *RIN* expression (Fig. S4). Other lines with large mutations spanning multiple CREs and DMRs all had suppressed *RIN*, i.e., ‘C5’, ‘A22’, ‘C3’, and ‘D25’; however, there was no correlation between the length of the deletion, and *RIN* expression. Notably, ‘A22’ is biallelic, where one allele has an 891-bp deletion and a 16-bp insertion in the *RIN* gene promoter, and the other allele has an insertion in a *RIN* exon. Both the T_0_ ‘A22’ and its heterozygote T_1_, ‘A22H’, which share the same genotype, had reduced *RIN* expression (Fig. 1F, S4).

### Fruit ripening phenotype

To assess if inducing mutations in *RIN* CREs affected fruit ripening, five mutant T_1_ lines were selected to evaluate fruit ripening speed, defined as the timeframe from the onset of ripening (Breaker stage) to Red Ripe. Multiple parameters were used to depict the dynamism of fruit ripening within each genotype ripening trends, including quartiles, mean values, i.e., average number of days fruit took to ripen, and a ripening index relative to WT (Fig. 1H). Strikingly, the ‘A22H’ mutant took an average of 11.7 days to ripen compared to 6 days in the WT, with a 15-fold higher ripening index. This ripening delay may be related to an allelic disruption in the *RIN* exon (Fig. 1J), which is also associated with reduced *RIN* expression (*P* < 0.01; Fig. S4). In contrast, other genotypes such as the small indel mutant ‘A26B’, the large deletion mutant ‘A26A’, and the heterozygous lines ‘A26H’ and ‘A83’, had ripening pattern and *RIN* expression at Red Ripe stage that were comparable to WT (*P* > 0.05; Fig. 1H, I, S4).

Fruit color was used as a visual marker for ripening progression; however, external color alone may not capture earlier molecular or physiological changes (Zhou et al., 2024). We further investigated whether promoter mutations affect fruit ripening and quality. Two homozygous mutants, ‘A26A’ and ‘A26B’, each carrying mutations at RIN binding site (Fig. 1J), were selected for further analysis. *RIN* gene expression was suppressed early, at day 1 (*P* < 0.05) and day 3 post-breaker but returned to WT-like levels by day 5 (Fig. 1K) in both genotypes.

Consistent with the gene expression pattern, both genotypes displayed a ‘delayed ripening’ phenotype (Fig. 1L). Assessment of two common fruit quality traits, total soluble solids (TSS) and firmness, revealed that ‘A26B’ fruit had significantly higher firmness (*P* < 0.05), while ‘A26A’ fruit had reduced TSS contents (*P* < 0.05). Together, these findings suggest that disruption of the *RIN* promoter can alter *RIN* transcription dynamics during early ripening and affect fruit quality traits, even without fully delaying the overall ripening timeline.

## CONCLUSION

We successfully generated various alleles, i.e., small indels, and large structural changes in the *RIN* gene promoter, resulting in diverse phenotypic outcomes. These include genotypes with extended ripening times (e.g., ‘A22H’), and others with subtle quality traits changes despite a normal ripening progression. Our work reveals a non-linear relationship between promoter mutations and gene expression levels, with certain types of mutations having a more significant effect than others. This study provides evidence that editing gene regulatory regions can create novel allelic resources varying incrementally in the expression of a ripening gene and alter fruit ripening physiology. Future work will focus on evaluating fruit traits from the mutants with differential *RIN* expression changes or distinct mutation types, as well as understanding if the mutant fruit will respond differently to postharvest stress.

## Declarations

### Ethics approval and consent to participate

The authors secured biological use authorization to generate the transgenic material described in this study.

### Consent for publication

Not applicable.

### Availability of data and material

The constructs and plant materials in this study are available in the lab of the corresponding author upon reasonable request. The data generated during this study are included in this published article and its supplementary information file.

### Competing interests

The authors declare that they have no known competing financial interests or personal relationships that influenced the work reported in this paper.

### Funding

Jiaqi Zhou: Horticulture and Agronomy Graduate Group Ph.D. Fellowship at UC Davis; Chiu-Ling Yang: Graduate Students Study Abroad Program grants from the Taiwan National Science and Technology Council (NSTC 112-2917-I-005-004); Diane M. Beckles: Hatch Project CA-D-PLS-2404-H and a UC ADVANCE Fellowship.

### Authors’ contributions

Jiaqi Zhou designed and performed the experiments, analyzed the data, wrote the initial and final drafts of the paper; Chiu-Ling Yang: analyzed data, generated figures, and edited manuscript drafts; Diane M. Beckles conceived, funded, and supervised the work, and edited initial and final drafts.

## Acknowledgements

We thank Hsin-Ya ‘Jasmine’ Hsiao (Provost’s Undergraduate Fellow) and Dr. Po-Kai Huang for their help with RNA isolation and qRT-PCR assessment of on-the-vine ripening fruit at different developmental stages. We are grateful to Dr. David M. Tricoli who offered critical advice on tomato transformation and regeneration. We thank UC Davis undergraduate interns - Daniel Leung, Roshmund Romero, Sheridan Chavira, Annika Uemura, Faizan Ashraf, Michael Muljadi, Juliana Amireh, Yuan Shu, Alexander Dominguez, Zhiying He, Sarah Raza, Sachi Nandedkar, Armaan Singh, and UC Davis Young Scholar Chloe Choi, who helped with tomato transformation and tissue culture, and genotyping.

## SUPPLEMENTARY MATERIALS

Supplementary Table 1. Guide-RNA sequences used.

Supplementary Table 2. Primers used for gene construct assembly.

Supplementary Table 3. Primers used for screening and genotyping transformants.

Supplementary Table 4. RT-qPCR Primer set for *RIN* gene expression.

Supplementary Figure 1. RIN gene promoter DNA methylation status.

Supplementary Figure 2. Schematic of the CRISPR/Cas9 multiplex constructs.

Supplementary Figure 3. Summary of small indels position and sequence in the T_0_ lines.

Supplementary Figure 4. Summary of small indels position and sequence in the T_0_ lines.

## Materials and Methods

**Table S1.**
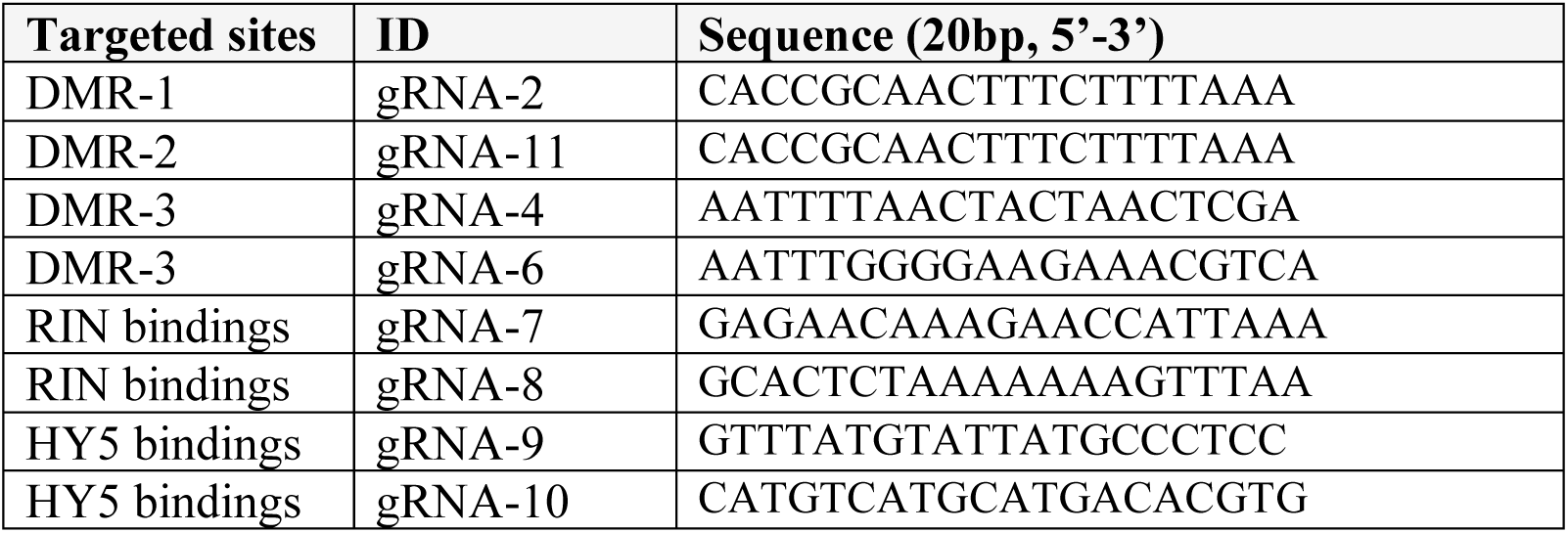
Guide-RNA sequences used.

**Table S2.**
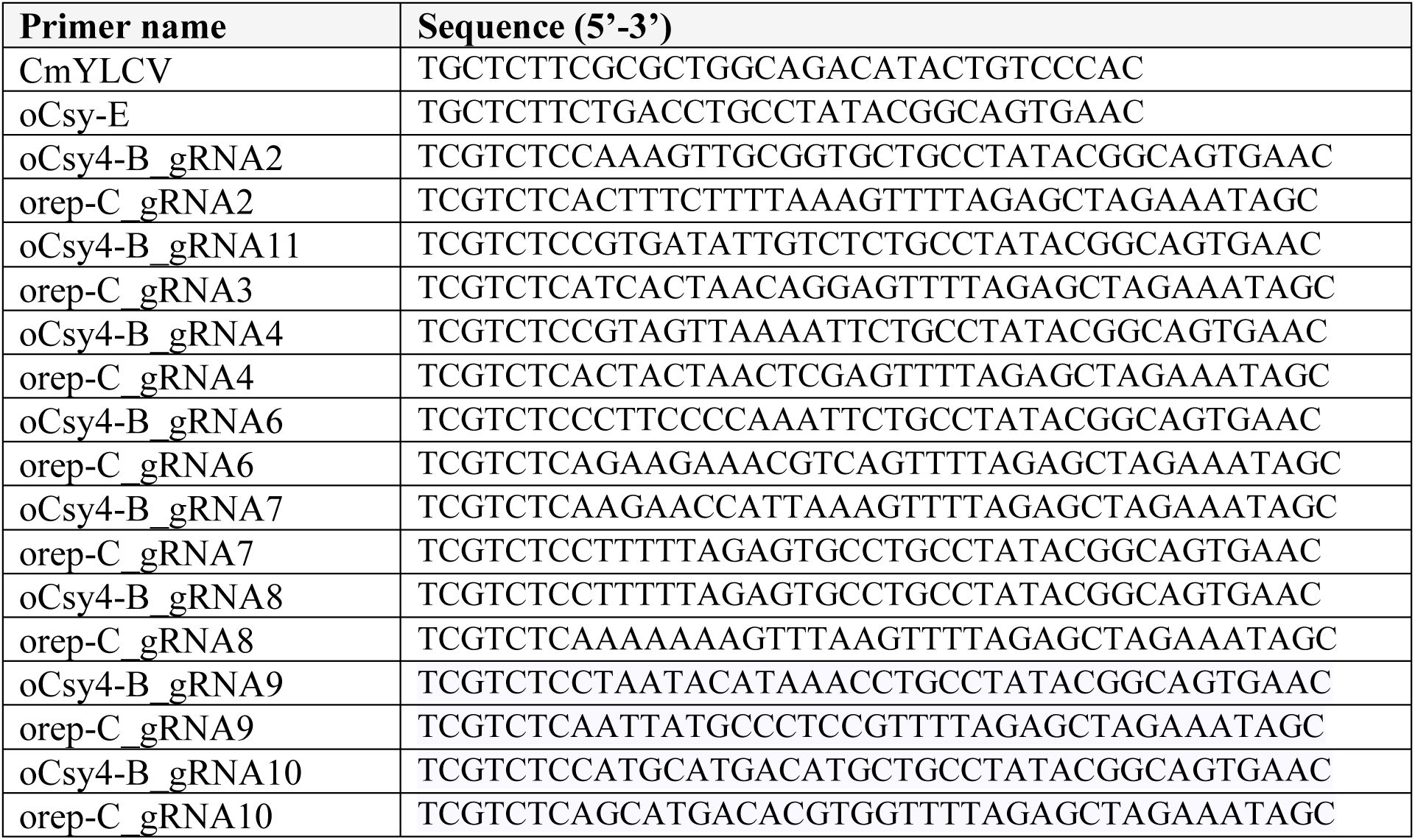
Primers used for gene construct assembly

**Table S3.**
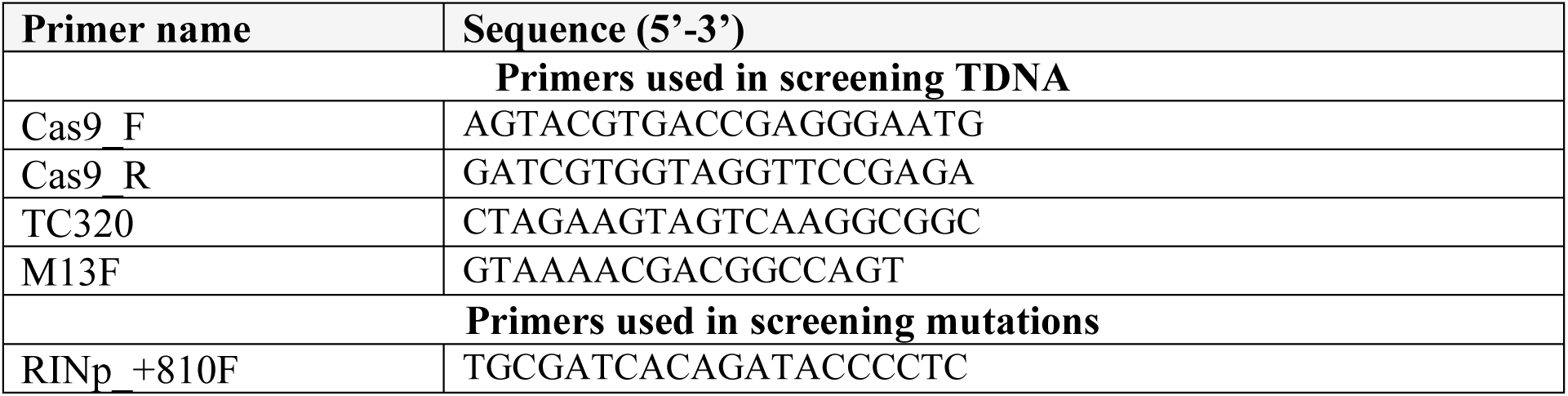
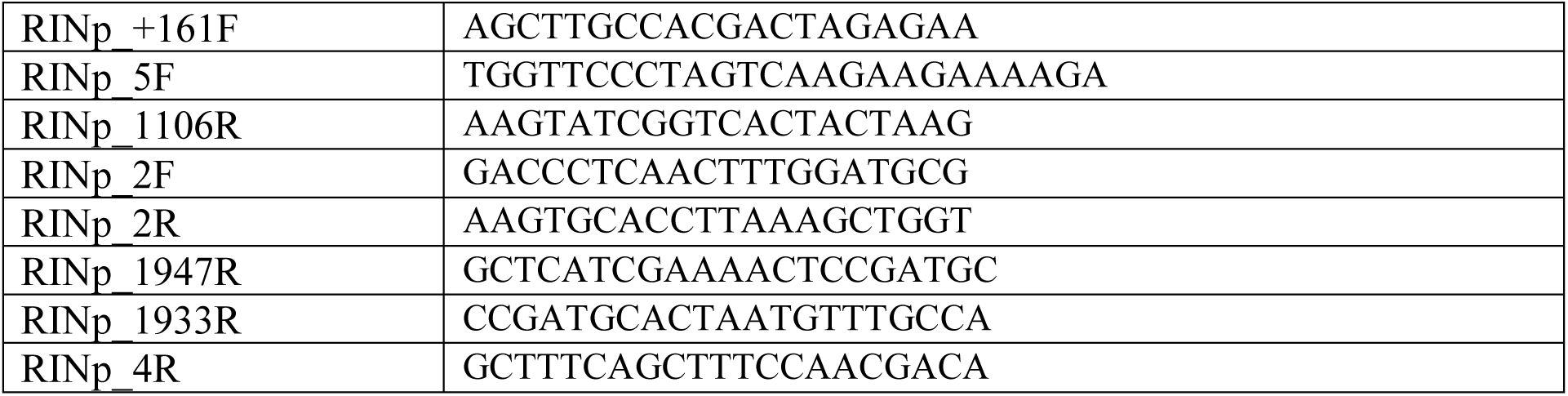
Primers used for screening and genotyping transformants

**Table S4.**
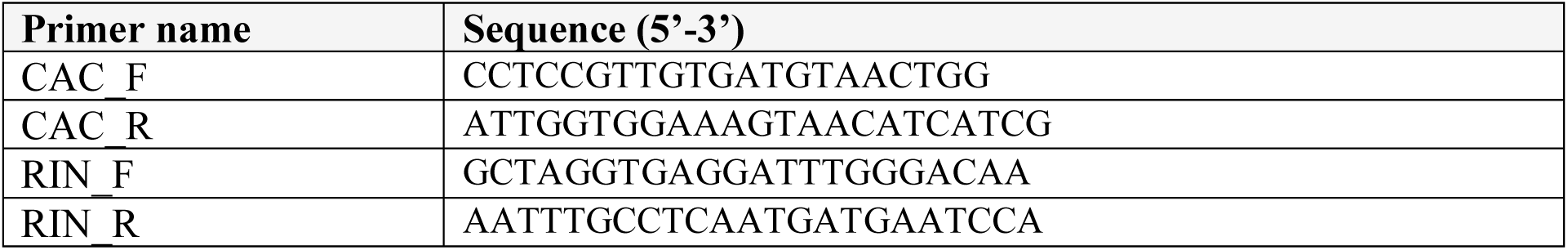
RT-qPCR Primer set for *RIN* gene expression (Ito et al., 2020)

## MATERIALS AND METHODS

**Figure S1.**
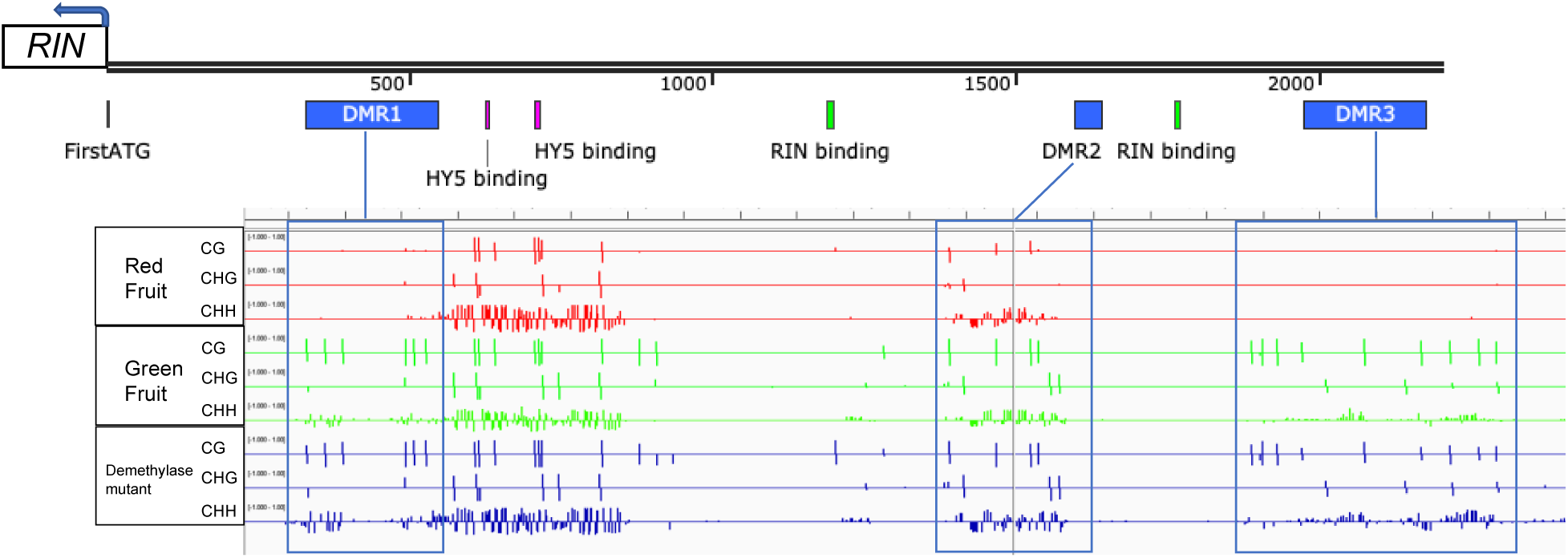
*RIN* gene promoter DNA methylation status. The image of raw DNA methylation sites was from Lang *et al*. (Lang et al., 2017). The single-base level DNA methylation for fruit tissue was shown by the CG, CHG and CHH contexts, respectively. The red panel is for tomato fruit at the red ripe stage, the green panel is the data for fruit before the onset of ripening, and the blue is for the Demethylase-*SlDML2* mutant. There are three DMRs identified and highlighted by the blue boxes. These regions are ripening-induced hypo-DMRs, because their DNA methylation levels drop during ripening. The sequence length of the DMRs is 218 bp, 45 bp and 201 bp, respectively.

**Figure S2.**
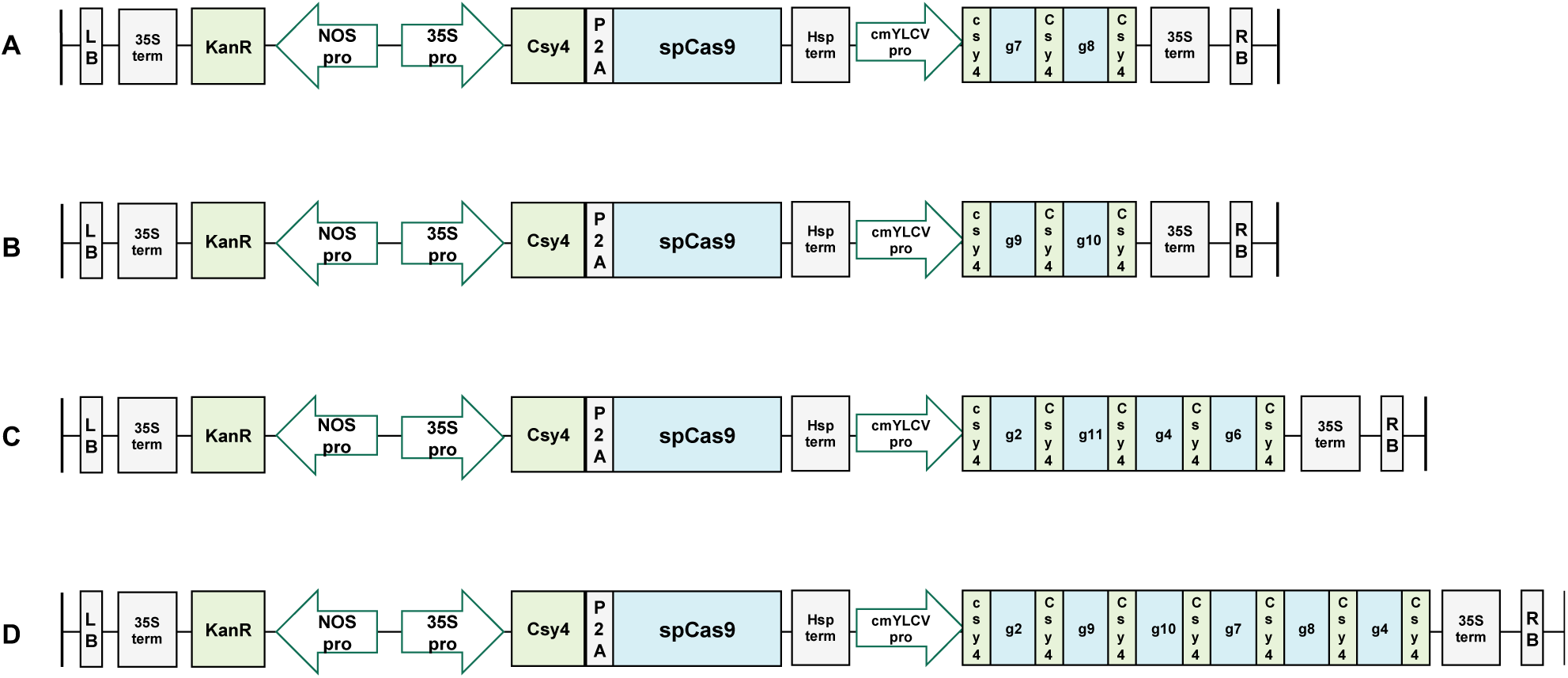
Schematic of the CRISPR/Cas9 multiplex constructs. A gene construct containing (**A**) Two gRNAs for editing the RIN self-binding sites. (**B**) Two gRNAs for editing the HY5 binding sites. (**C**) Four gRNAs for editing the three hypo-DMRs that are described above. (**D**) Six gRNAs for the simultaneous targeting of the CREs to which RIN and HY5 bind, and the DMRs.

**Figure S3.**
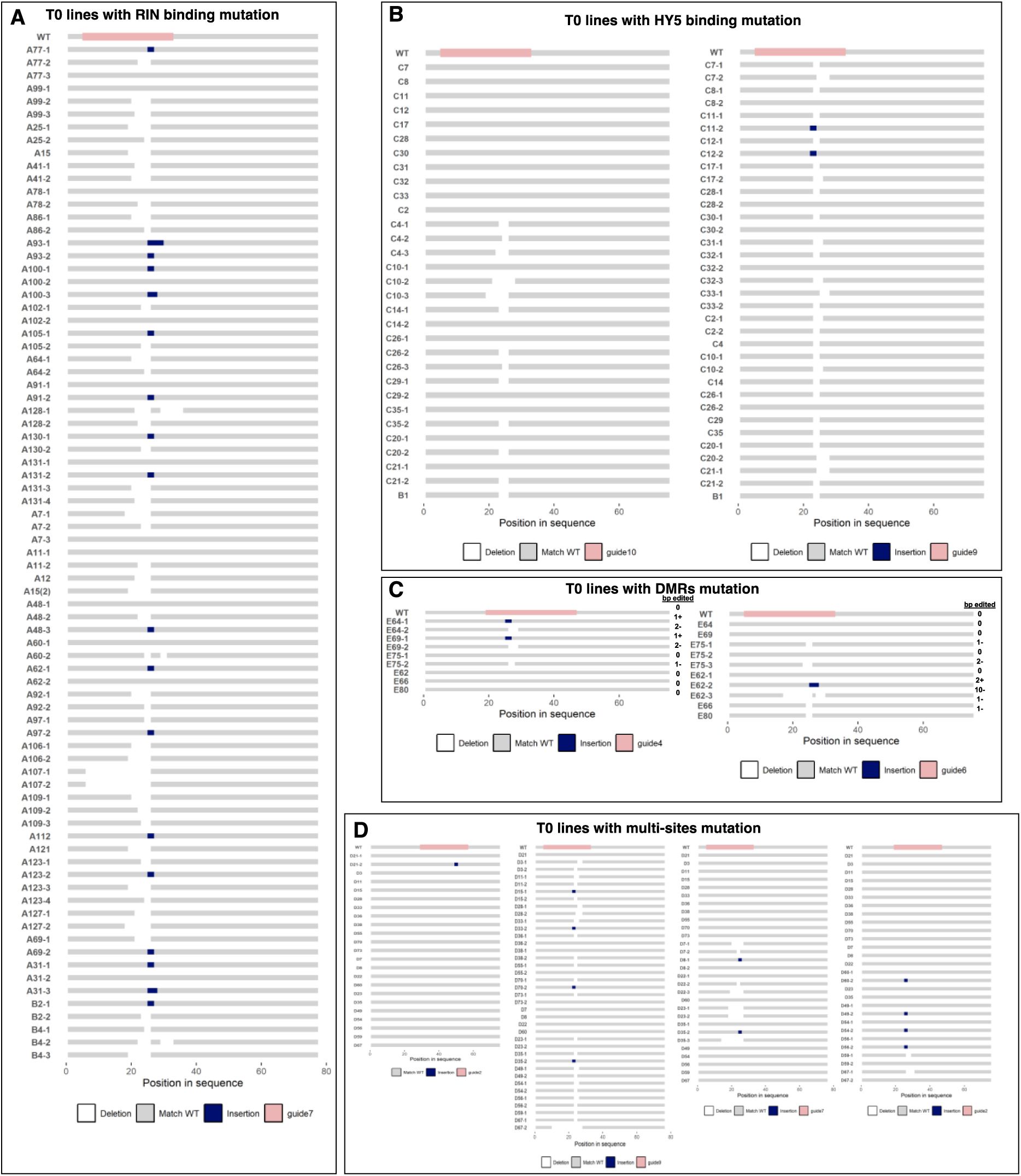
Summary of the small indels (insertion or deletion) position and sequence in the T_0_ lines. Mutations were presented at (**A**) RIN binding site, (**B**) HY5 binding site, (**C**) DMRs region, (**D**) combination of RIN binding site, HY5 binding site, and DMR region. There are multiple alleles in some heterozygous or chimeric T_0_, represented by an addition of ‘ -1, -2, or - 3’ after the T_0_ line’s ID. The grey bars indicate the same sequence as WT, the white areas indicate ‘deletion’ regions, and the blue bars indicate those with ‘insertions.’ The pink bar at the top of each column indicates the gRNA targeting region (20 bp).

**Figure S4.**
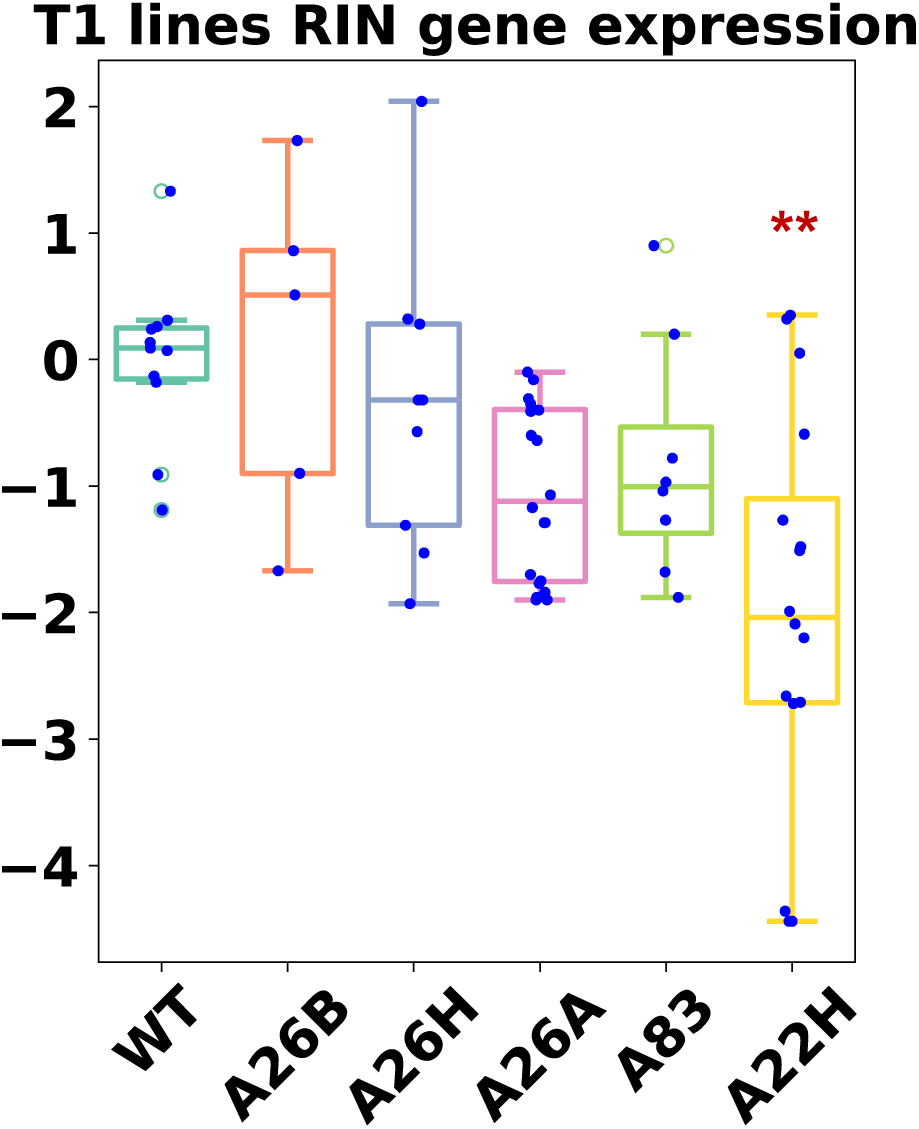
Relative expression of *RIN* in red ripe fruit of WT and T_1_ edited lines. The y-axis indicates the log_2_ fold change relative to WT. Compared to WT, *P* < 0.01 is indicated by ‘**’, according to the one-way ANOVA test with Tukey’s multigroup correction.

### 1. RIN gene promoter analysis

The *RIN* promoter region is defined as the 2-3 kb region upstream of the *RIN* gene (NCBI ID: 543708) translation start site. The sequence was accessed through NCBI (JAAXDC010000005.1:5415385-5418385 *Solanum lycopersicum* cultivar Micro-Tom chromosome 5, whole genome shotgun sequence).

The sequence was used as the input for identifying potential cis-regulatory regions (CREs) analysis in PlantCare (Lescot et al., 2002). The two identified CArG motifs to which RIN binds were identified according to (Bemer et al., 2012; Fujisawa et al., 2013). The DNA methylation status of the RIN gene promoter in ‘Micro-Tom’ was downloaded from (https://www.ncbi.nlm.nih.gov/geo/query/acc.cgi?acc=GSE94903) (Lang et al., 2017). The downloaded sequence data with DNA methylation in three contexts, i.e., CG, CHH and, CHG were imported to the Integrative genomics viewer (IGV) (Robinson et al., 2011). Comparing the DNA methylation levels in wild-type fruit at different ripening stages, the ripening-induced differential methylated regions (DMRs) were identified.

### 2. Gene construct assembly

The gRNAs targeting the three ripening-induced hypo-DMRs in the *RIN* promoter, two RIN self-binding sites (CArG motif), and HY5 binding sites were designed using CRISPOR (Concordet & Haeussler, 2018) and CRISPR-P (Lei et al., 2014). To avoid ‘off-targeting’, the gRNA sequences were selected based on the off-target mismatches score, followed by BLAST against the Micro-Tom genome through NCBI. The CRISPR/Cas9 constructs were generated using the pDIRECT_22C (Plasmid #91135, Addgene) backbone through Golden Gate cloning (Čermák et al., 2017). Some constructs carried gRNAs targeting a single type of site, while others carried multiple gRNAs to target different types of sites simultaneously. Assembled constructs were transformed into *E.coli* strain DH5-alpha. Plasmid DNA was isolated using a miniprep kit (Qiagen, Valencia, CA) and whole plasmid sequencing was done using the service offered by Plasmidsaurus (Eugene, OR, USA).

### 3. Plant transformation and tissue culture

Sequenced plasmids were transformed into *Agrobacterium tumefaciens* strain GV3101 using the heat-shock method (Goldbio, MO, USA). *A. tumefaciens-*mediated plant transformation was done as described in Albornoz *et al*. (Albornoz et al., 2023). Tomato tissue culture were performed under the guidance of the Plant Transformation Facility at UC Davis. The regenerated tomato plantlets with well-developed roots and shoots were transplanted into soil and hardened under controlled environmental conditions for two weeks. The adapted plants were transferred into the greenhouse at Davis, CA.

### 4. Plant growth and fruit sampling

All mutants and WT plants were grown in the greenhouse at UC Davis (Davis, CA, USA) using a completely random design (CRD). The UC ‘Mix C’ soil was placed in a 6.5-inch pot and watered (with liquid fertilizer) twice a day with a 3 min irrigation cycle. The greenhouse conditions were maintained at 25°C, with a 15-h light and 9-h dark cycle each day. The fruit were harvested and washed in 0.25% (v/v) sodium hypochlorite for three minutes, followed by rinsing with water and gently blotted until dry with paper towels. Fruit pericarps were sampled, frozen in liquid nitrogen and stored at -70°C for RNA isolation.

### 5. Mutation detection

The following genotyping standard procedures were applied for hundreds of regenerated first-generation (T_0_) plantlets. (1) Genomic DNA was extracted from young tomato leaves using a modified CTAB method (Zhou et al., 2021). (2) Genomic DNA quality was tested by a Nanodrop and a standard PCR amplification for a housekeeping gene *ACT7* using AmpliTaq (Applied biosystem, USA). (3) The genomic DNAs that passed the quality checks were used for screening T-DNA insertion by amplifying the Cas9 fragment of the construct. (4) Genomic DNA from individual lines with T-DNA insertions were used to amplify the gRNAs regions in the *RIN* promoter sequence, using flanking primers. (5) Direct Sanger sequencing was used to analyze the purified amplicons. The mutation type, i.e., homozygous, biallelic, heterozygous and chimeric (multiple mutations) was detected by NCBI blastn, TIDE (Brinkman et al., 2018) and ICE Synthego (Conant et al., 2022) together. Plants in the T_1_ generation were genotyped by PCR-amplifying the entire region targeted by the gRNAs, and subjecting the purified fragments to Sanger sequencing to detect small indels. To generate the aligned sequence figures, the sequence traces were first aligned in MEGA X software (Kumar et al., 2018), illustrated by the seqvisr and the ggmsa package (Charif & Lobry, 2007; Zhou et al., 2022) under the R environment.

### 6. RNA isolation and RT-qPCR

Fruit total RNA was isolated from around 100 mg fruit power using a Trizol-based protocol (Invitrogen, Thermofisher, USA). RNA quality and integrity were assessed by microvolume spectrophotometer and 0.8% (w/v) agarose gel electrophoresis. Around 500 ng total RNA was used as the input for the High-capacity Reverse Transcription kit (Thermofisher). The cDNA libraries were quality checked using a standard PCR (Amplitaq; Applied Biosystems). The RT-qPCR were performed according to Zhou *et al*. (Zhou et al., 2021), using the *SlCAC* gene as the internal control as reported (Ito et al., 2020).

### 7. Fruit ripening speed assessment

The five T_1_ generation line assessed, included ‘A26A’, ‘A26B’, ‘A26H’, ‘A22H’, ‘A83’, and were compared to the WT. Ripening time was recorded as the duration between Breaker and Red Ripe fruit ripened on-the-vine. There were 114 WT fruit, 75 fruit for ‘A22H’, 57 fruit for ‘A26A’, 28 fruit for ‘A26H’, 24 fruit for ‘A26B’, and 32 fruit for ‘A83’, harvested from at least five tomato plants per genotype in this assay. All plants were grown simultaneously under the same conditions in the greenhouse at UC Davis in Summer 2023, and the data were recorded between 9 am -10 am daily.

### 8. Fruit quality parameters assessment

Fruit were harvested at Breaker, and stored at 20°C with 67% relative humidity under dark conditions for one week, to evaluate fruit quality. Fruit firmness was assessed using a Texture Analyzer (TA. XT Plus, Texture 147 Technologies, Scarsdale, NY). The force was recorded when compressing the equatorial region of the whole fruit for 5 mm using a 2 mm puncture flat stainless probe. The parameters were set as 2 mm/s for the pre-test speed, 1 mm/s for the test speed, followed by a 5 mm/s post speed. The average value was determined at three points for each fruit. Total Soluble Solids (TSS) was assayed by placing fruit juice in a refractometer at room temperature (Hanna Instruments, USA) (Zhou et al., 2021). The TA was represented as grams of citric acid per 100 g of fresh tomato weight.

### 9. Statistical analysis

A completely randomized design (CRD) was applied, in which treatment levels included genotype and ripening stages. Significant differences (*P* < 0.05) were determined by ANOVA or Student’s *t*-test in the R-environment.

## Notes

### Competing Interest Statement

The authors have declared no competing interest.

